# flodl-bio: First Rust data, transformers and model layer for entire deep learning on biological sequences

**DOI:** 10.64898/2026.10.21.760383

**Authors:** Gaurav Sablok, Fabrice de Stefanis

## Abstract

Python ecosystem dominates the deep learning framework. flodl-bio is a Rust crate that extends flodl, a libtorch-backed deep-learning framework, with deep learning on biological sequences. To our knowledge, flodl-bio is the first Rust crate that provides a comprehensive and practical deep learning for the biological applications, which includes the alphabet-aware sequence encoding, including one-hot and k-mer representations, padded sequences, and channels-first batches. It also includes several FASTA parsing and graph construction utilities for adjacency matrices, GCN-normalised adjacency matrices, graph Laplacians, residue contact maps, and de Bruijn graphs. Among the added functionalities, it includes AUROC and AUPRC with their associated curves; classification metrics, and the export of learned convolutional filters as MEME motifs and seeded splits. Deep biological learning includes a multi-width motif CNN, a dilated residual CNN, bidirectional LSTM and GRU architectures, and transformer encoder. On the five human transcription-factor-binding datasets in the GUE benchmark, a motif CNN with approximately 10,000 parameters, trained from scratch, outperforms all published CNN baselines. It also superseded the 2.5-billion-parameter Nucleotide Transformer on two datasets and DNABERT-2 on one. Given a fixed hardware configuration and software stack, experiments are reproducible at the bit level. See Supplementary Table S2 of Supplementary Material.

**Availability and implementation:** flodl-bio is written in Rust and released under the MIT licence at https://github.com/sablokg/flodl-bio. The core builds with stable Rust and has no dependencies. The model tier is enabled with the flodl Cargo feature (cuda for NVIDIA GPUs) and requires libtorch and Rust 1.91 or later. The evaluation harness is in the repository’s experiments directory, and the complete run outputs are archived at https://doi.org/10.5281/zenodo.23321850.

## 1 Introduction

Convolutional, recurrent, attention-based, and graph neural networks have become standard tools for regulatory genomics, protein modelling, and variant-effect prediction (Alipanahi et al., 2015; Zhou and Troyanskaya, 2015; Quang and Xie, 2016; Avsec et al., 2021). Most existing tooling is built on Python-based frameworks such as PyTorch (Paszke et al., 2019). Rust offers memory safety, predictable performance, single-binary deployment, and seamless integration with the Rust bioinformatics ecosystem (Köster, 2016), making it an attractive choice for production pipelines and tools embedded in sequence-processing software.

flodl is a Rust deep-learning framework built on libtorch that provides tensors, automatic differentiation, a broad range of layers, loss functions, optimizers, and learning-rate schedulers (flodl, 2026). However, it lacks domain-specific data-handling utilities and reference architectures for biological applications. Consequently, users must implement their own sequence encoders, graph-construction routines, batching strategies, and evaluation pipelines. flodl-bio addresses this gap. To our knowledge, it is the first Rust package to support an end-to-end deep-learning workflow for biological sequences, from data encoding to model training, including Transformer architectures. Existing Rust crates, such as DeepBioP (2024), focus on preparing biological data for use with Python-based frameworks and do not provide an integrated training and optimization pipeline.

Regulatory genomics task involves the training of the pre-trained DNA language models. It has been shown previously that the supervised models trained from baseline using one-hot encoded DNA sequences and Convolutional Neural Networks (CNNs) are competitive with the ones trained on the frozen representations (Tang et al., 2025) or embeddings (Marin et al., 2024) using pre-trained language models (Marin et al., 2024; Tang et al., 2025). It has been shown that the transcription-factor-binding tasks in the GUE benchmark as per the previously published CNN baselines substantially underperform DNABERT-2 (Zhou et al., 2024; Karpinsky et al., 2026). We used the evaluation framework provided by flodl-bio to revisit this comparison using compact models and a reproducible experimental protocol (Section 3.2).

## 2 Features

flodl-bio is organised in two tiers (Supplementary Table S1). The **core** has no dependency on libtorch and covers data preparation. The **model** tier, behind the flodl cargo feature, covers architectures and training.

### 2.1 Sequence encoding

Sequences are mapped to one-hot matrices of shape [length, alphabet]. Unknown symbols (for example N or X) either raise an error (strict mode) or become all-zero rows (lenient mode). Batches are zero-padded or truncated to a fixed length and accompanied by a mask. A channels-first transform produces the [batch, alphabet, length] layout used by one-dimensional convolutions. k-mer count and frequency vectors of length A^k support non-neural baselines and hybrid models.

### 2.2 Graphs

Graphs are stored as weighted edge lists and converted into dense row-major adjacency matrices. The GCN propagation matrix of Kipf and Welling (2017), the combinatorial Laplacian and an additive attention mask for graph attention (Veličković *et* al., 2018) are generated directly. Residue contact graphs are built from a distance matrix with a distance cutoff and a minimum sequence separation, and de Bruijn graphs are built from reads over (k-1)-mer nodes with multiplicity-weighted edges.

### 2.3 Models

MotifCnn utilizes parallel one-dimensional convolutional filters with varying widths, which are followed by ReLU activations and global max pooling, in accordance with the framework established by DeepBind (Alipanahi et al., 2015). This approach also allows for the interpretation of learned filters as sequence motifs. For each filter, input windows that achieve activations greater than half of the maximum activation are gathered and averaged to form a position frequency matrix (Kelley et al., 2016). The motifs derived from this process are then exported in MEME format, facilitating their comparison with motif databases through the use of TOMTOM (Gupta et al., 2007).

The DilatedResCnn architecture incorporates residual convolutional blocks with dilation rates that increase exponentially. The BiRnnClassifier integrates a forward recurrent network with an additional network that processes the reversed input sequence, thus offering bidirectional context, even though the LSTM and GRU layers are inherently unidirectional. The TransformerClassifier employs a pre-LayerNorm encoder featuring multi-head self-attention (Vaswani et al., 2017), sinusoidal positional encodings, and optional attention masking. The FlatMlp serves as a baseline multilayer perceptron, functioning on flattened one-hot-encoded inputs. We evaluated the convolutional-stem architecture in Section 3.2, which relies on Sequential, Residual, ConcatBranches, Lambda API. Complete runnable examples are mentioned in the README of flodl-bio, which can be accessed at https://github.com/sablokg/flodl-bio

### 2.4 Examples and reproducibility

flodl-bio includes four data-layer programs that run without libtorch, comprising three introductory walkthroughs and a throughput benchmark, and seven model-layer programs covering four architecture walkthroughs, a model-composition example, a model-throughput benchmark, and a FASTA-based evaluation harness that compares all supported sequence models for transcription-factor-binding prediction. A separate experiments crate reproduces the evaluation described in Section 3.2 with a single command. For each run, it generates a flodl training dashboard, per-epoch metrics, predictions, ROC and precision–recall curves, and a record of the configuration, hardware, and flodl-bio commit. For MotifCnn, the outputs additionally include the learned motifs in MEME format.

## 3 Evaluation

### 3.1 Data-layer throughput (measured)

Run conditions (bench_core, release build, commit 2cf9199): one core of an AMD Ryzen 9 9900X, best of five runs, median over three invocations, all within a 10% margin. Supplementary Table S2 lists one-hot encoding running at 177 Mbases/s, padded batch + channels-first at 97 Mbases/s, with the reverse complement reaching 6.2 Gbases/s. The GCN-normalised adjacency for a 500-node graph takes about 0.22 ms. The de Bruijn graph comes last with 4.3 Mbases/s due to one allocation per (k-1)-mer key, likely reducible with 2-bit key packing.

### 3.2 Transcription-factor binding on GUE

All sequence models were evaluated on the human TF-binding task in the GUE benchmark (Zhou *et al*., 2024), including five datasets from ENCODE ChIP-seq (ENCODE Project Consortium, 2012). They cover RNA polymerase II (two cell types, HUVEC and progenitor fibroblasts), PAX5, KAP1 (TRIM28) and MXI1 (Supplementary Note S2). We used 101-bp peak-centred sequences with GC-matched negative examples. Every model was trained from scratch for 20 epochs, using Adam with a batch size of 64, gradient-norm clipping at 1.0 and a constant learning rate (ranging from 5×10^-4^ to 2×10^-3^ depending on the architecture). No early stopping and no checkpoint selection were applied. The development (dev) split was only used to monitor training curves. Training splits range from 19,000 to 32,378 sequences, with dev and test splits of 1,000 sequences each. Table 3 displays test-set Matthews correlation coefficients (MCC), the GUE evaluation metric, with means and sample standard deviations over five random seeds (each setting initialisation, dropout and batch order).

**Table 3.**
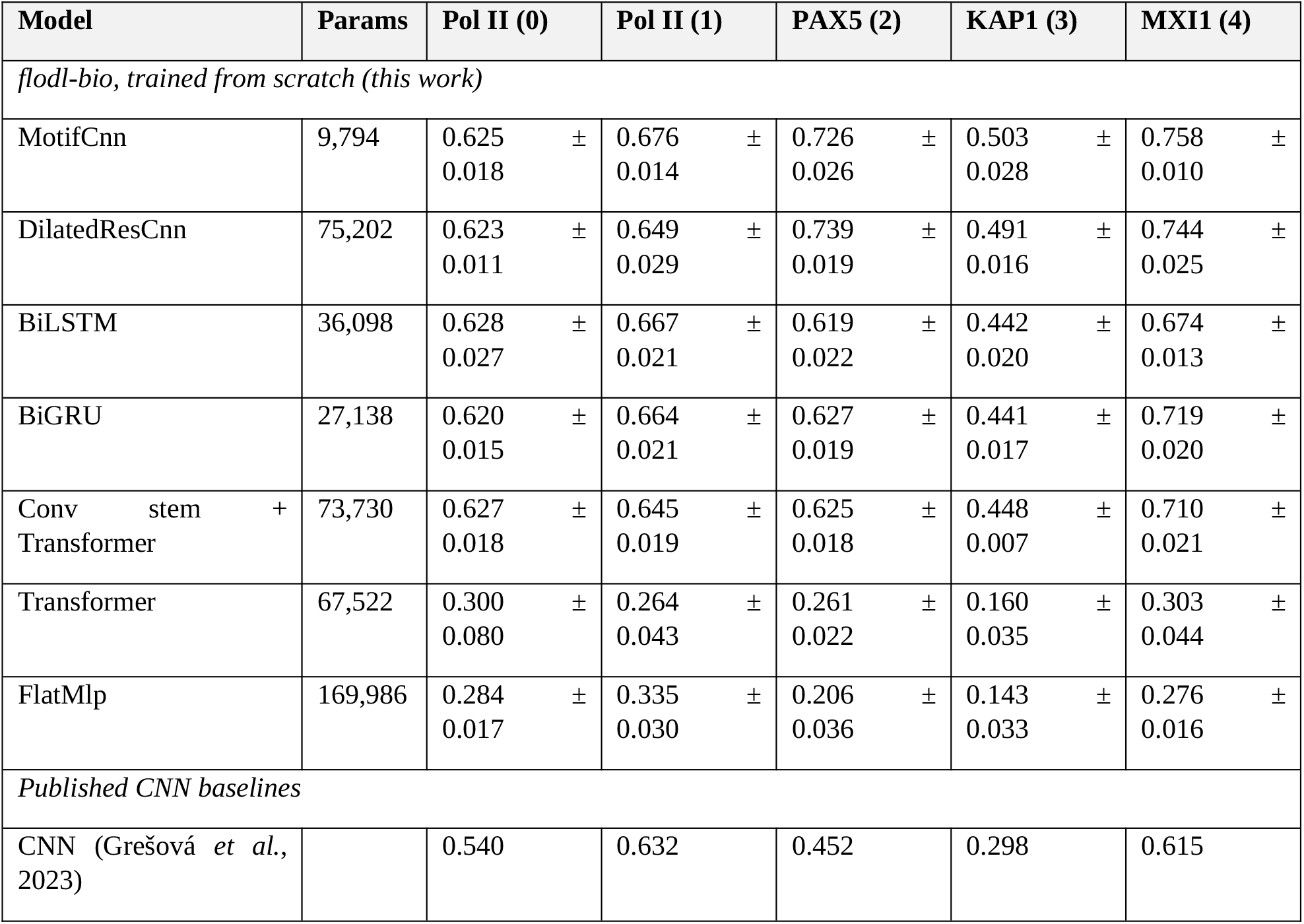
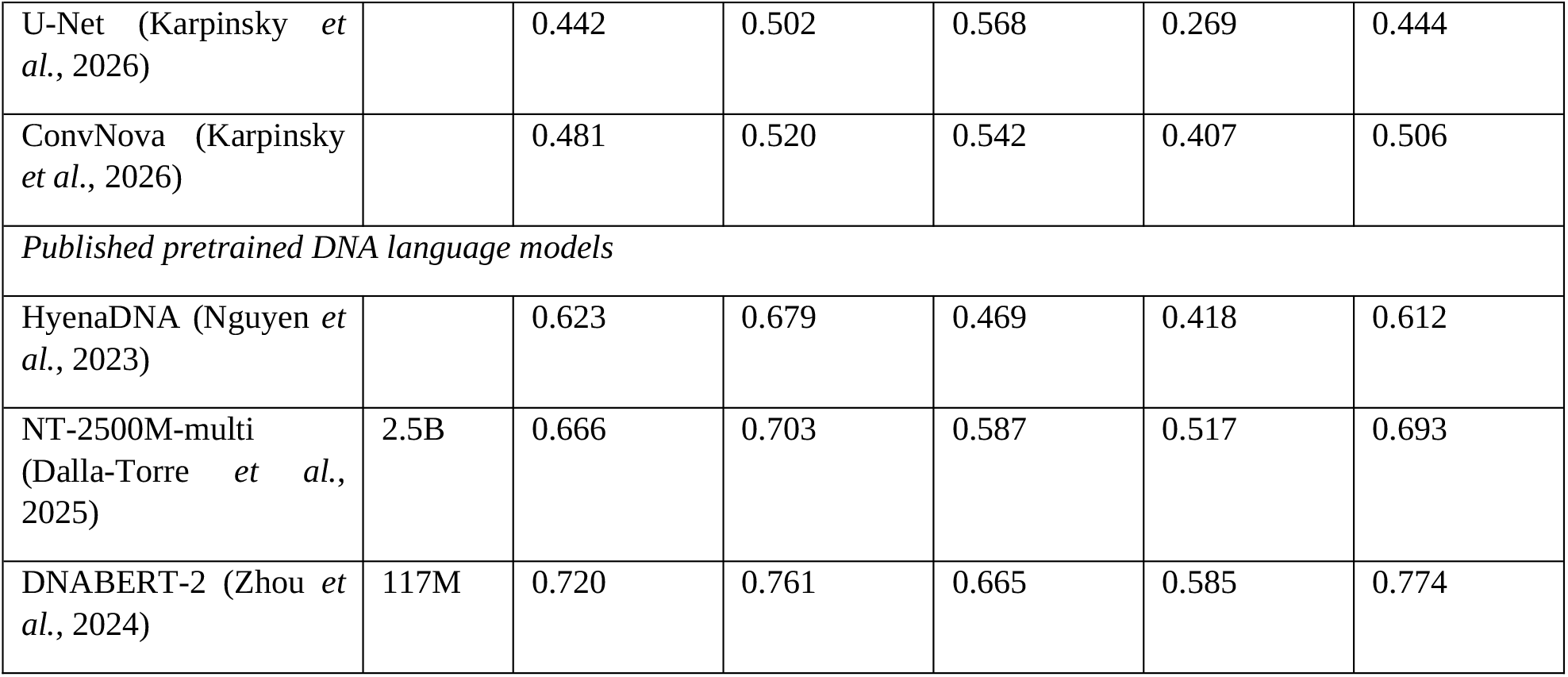
MCC from the GUE human TF-binding test sets. The published values are as reported by Zhou *et al*. (2024, Tables 6 and 9) and Karpinsky *et al*. (2026, Table 12). The best of three pre-training regimes on each dataset is used for U-Net and ConvNova. flodl-bio models are means ± sd over 5 seeds on an NVIDIA RTX 5060 Ti.

Interestingly, the smallest model (MotifCnn) has the best results among flodl-bio models on three datasets and stays within 0.013 MCC of the best one for the other two. It stays ahead of every published CNN baseline for all five datasets (0.04 to 0.16 MCC improvement). This comparison is conservative, at least against the CNN baseline of Zhou *et al*. (2024), who selected the checkpoint with the lowest validation loss while we only used the model after the final (20th) epoch for evaluation. MotifCnn, despite its size, also surpasses the 2.5-billion-parameter Nucleotide Transformer (Dalla-Torre *et al*., 2025) on PAX5 and MXI1. DNABERT-2 is behind on PAX5, but keeps a 0.08 to 0.10 lead on the two RNA polymerase II (Pol II) datasets and KAP1, and 0.02 on MXI1. As training protocols differ across studies, it is not possible to attribute any observed gains to a single factor. We observed an unsurprising underfitting of the standard Transformer encoder at these dataset sizes, and found that adding a single convolutional stem (using the composition containers) brought its performance in line with the recurrent architectures. Performance fits the biological complexity with PAX5 and MXI1 having strong sequence-motif signals (Supplementary Table S4) while KAP1 remains the most challenging, as it lacks a DNA-binding domain of its own. Pol II sits in between with its diffuse features.

The GPU is a consumer NVIDIA RTX 5060 Ti, the CPU is an AMD Ryzen 9 9900X and all training ran with libtorch 2.10, CUDA 12.8 and flodl-bio at commit b6ce8b4. MotifCnn trained at 0.26 s per epoch on GPU, and 0.56 s per epoch on CPU (32,378 sequences), with around 25 MB of GPU memory allocated to tensors, or 200 MB RAM (including libtorch overhead) when trained on CPU. The complete evaluation used 175 runs and lasted about 30 min on the GPU and 4 h 20 min on the CPU. A complete re-run of the 175 runs on each device reproduced identical predictions, curves and per-epoch metrics, demonstrating bit-for-bit reproducibility under the same hardware and software. GPU and CPU execution followed different numerical paths and gave different per-seed results, but their means stayed within seed variability (Supplementary Table S3). Three out of 70 MCC and AUROC comparisons fell outside two standard errors, which stays close to the 3.5 exceedances expected by chance under a normal approximation.

**Figure 1:**
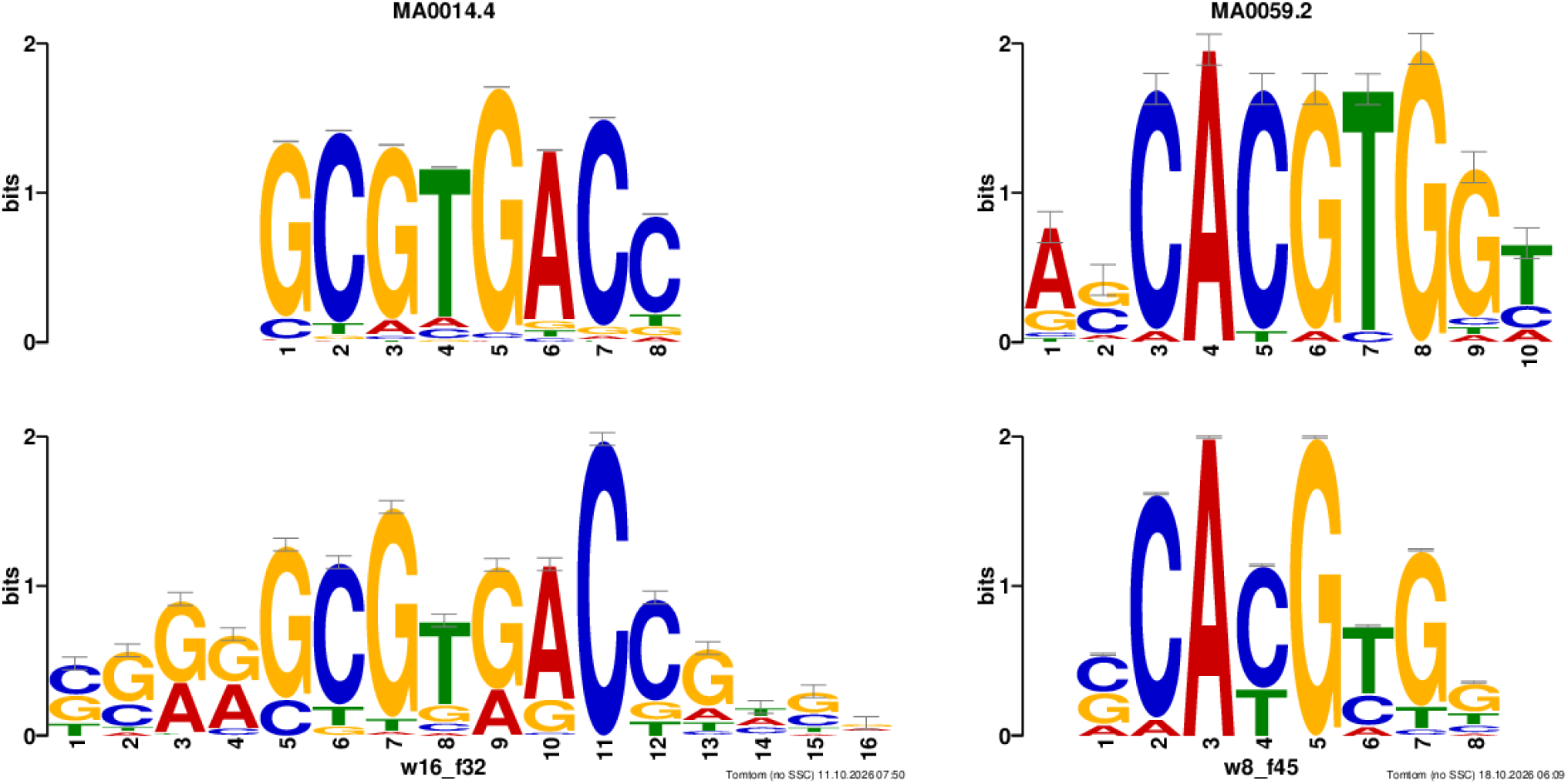
learned MotifCnn filters for PAX5 and MXI1 beside their best TOMTOM matches in JASPAR. w8_f45 learned on the MXI1 with MA0059.2 (MAX::MYC), q = 0.00013 as the best match and w16_f32 learned on the PAX5 with MA0014.4 (PAX5), q = 0.011 as the best match. For TOMTOM analysis, seed 1 was used and then searched against the JASPAR 2026 CORE vertebrates non-redundant collection. The five-seed results are in Supplementary Table S4.

## 4 Discussion and limitations

flodl-bio has been developed to reduce the effort of developing biological machine and deep learning models in Rust. We evaluated on GUE and it has been shown that the compact, supervised models trained from scratch remain strong baselines for transcription-factor-binding prediction, consistent with evaluations of pretrained DNA language models on other tasks (Marin et al., 2024; Tang et al., 2025). flodl-bio has several limitations such as the evaluation covers a single task family with no hyperparameter search and within one benchmark. Additionally, dense adjacency matrices have been used to scale graph instead of the sparse matrices, which can scale large graphs and are not yet included in this release version. The training loop does not restore the epochs weights which are best-performing and bidirectional recurrent architectures are implemented using a second network operating on reversed sequences . flodl-bio does not include pretrained genomic or protein language models. Two- and three-dimensional convolutional architectures can be assembled through the composition API but are not provided as dedicated reference implementations. Finally, because the modelling layer depends on flodl, the stability of flodl-bio’s interface is constrained by the stability of its underlying framework.

## Author contributions

G.S.: coded flodl-bio, ran TOMTOM analysis, wrote all test runs and wrote original draft, review and editing draft. F.d.S.: Software, Resources (computing), Investigation (running the trainings), Validation (reproducibility re-runs, GPU tests and GPU/CPU agreement checks), Formal analysis (multi-seed statistics, sd, CPU vs GPU significance count, and the five-seed TOMTOM run and analysis), Methodology (GUE protocol, fixed epochs, no checkpoint selection and seed count), Writing (original draft of Section 3, review and editing).

## Use of AI tools

G.S. used ChatGPT (OpenAI) to clarify and check algorithms and methods implemented during development, grammar correction and language corrections. F.d.S. used Claude (Anthropic), through Claude Code, to assist with software development, evaluation runs, analysis, literature search, grammar, spelling and consistency checks of this manuscript (Supplementary Note S1).

