## Supplementary Material (Tables S1-S4, Notes S1-S2) for "flodl-bio: First Rust data, transformers and model layer for entire deep learning on biological sequences"

**Table S1.** The main components of flodl-bio are separated between data preparation (no dependencies), and models (architecture and training) which rely on flodl (depends on libtorch and Cargo features).

| Module | Contents |
|---|---|
| alphabet | DNA, RNA and 20-residue protein alphabets (case-insensitive); reverse complement; GC content |
| encode | strict and lenient one-hot encoding; class-label one-hot; padded batches with masks; channels-first layout for convolutions; k-mer counts and frequencies; sinusoidal positional encodings |
| io | FASTA parsing; centre-cropping or padding to a fixed length |
| graph | dense adjacency; GCN-normalised and row-normalised adjacency; Laplacian; additive attention mask; edge index; connected components; contact-map graphs from 3-D coordinates; chain graphs; de Bruijn graphs |
| data | in-memory datasets; sequence-to-dataset construction; seeded shuffling and random train/validation split; accuracy, confusion matrix, macro-F1, MCC, AUROC, AUPRC; ROC and precision-recall curves; synthetic motif and graph tasks |
| motif | position frequency matrices from convolutional filter activations; background letter frequencies; MEME export for TOMTOM |
| models | MotifCnn, DilatedResCnn, BiRnnClassifier (LSTM or GRU), TransformerClassifier, FlatMlp; composition containers (Sequential, Residual, ConcatBranches, Lambda) |
| train | fit and fit_with (per-epoch callback; gradient clipping, learning-rate decay, early stopping), evaluate, predict_proba, report |
| prelude | one import that re-exports all of flodl together with the items above |

**Table S2.** Data-layer throughput, measured as the best of five runs, median over 3 invocations, on a single core of the AMD Ryzen 9 9900X CPU used.

| Operation | Input | Time (ms) | Throughput |
|---|---|---|---|
| One-hot encoding | 2,000 × 1,000 bp | 11.3 | 177 Mbases/s |
| Padded batch + channels-first | 2,000 × 1,000 bp | 20.7 | 97 Mbases/s |
| k-mer counts (k = 6) | 2 Mbp | 39.9 | 50 Mbases/s |
| Reverse complement | 2 Mbp | 0.33 | 6,150 Mbases/s |
| GCN-normalised adjacency | 500 nodes | 0.22 | 1,160 Mcells/s |
| Laplacian | 500 nodes | 0.24 | 1,050 Mcells/s |
| Contact graph from coordinates | 500 residues | 0.12 | 2,140 Mcells/s |
| de Bruijn graph (k = 21) | 200 reads × 1,000 bp | 46.7 | 4.3 Mbases/s |
| AUROC | 10^6 scores | 27.5 | 36 Mscores/s |

| Operation | Input | Time (ms) | Throughput |
|---|---|---|---|
| Average precision (AUPRC) | 10^6 scores | 25.7 | 39 Mscores/s |
| Dataset split + batch gather | 2,000 × 1,000 bp | 1.87 | 1,070 Mbases/s |

**Table S3.** MCC and AUROC are mean ± sd over 5 seeds, using an AMD Ryzen 9 9900X CPU. Test-set results follow the same protocol as Table 3.

| **Model** | **Pol II (0)** | **Pol II (1)** | **PAX5 (2)** | **KAP1 (3)** | **MXI1 (4)** |
|---|---|---|---|---|---|
| *MCC* | | | | | |
| MotifCnn | 0.632 ± 0.007 | 0.667 ± 0.021 | 0.707 ± 0.012 | 0.506 ± 0.012 | 0.751 ± 0.016 |
| DilatedResCnn | 0.613 ± 0.023 | 0.639 ± 0.018 | 0.739 ± 0.026 | 0.504 ± 0.023 | 0.741 ± 0.014 |
| BiLSTM | 0.643 ± 0.013 | 0.652 ± 0.007 | 0.611 ± 0.016 | 0.438 ± 0.022 | 0.690 ± 0.017 |
| BiGRU | 0.618 ± 0.016 | 0.655 ± 0.022 | 0.617 ± 0.023 | 0.435 ± 0.035 | 0.701 ± 0.014 |
| Conv stem + Transformer | 0.641 ± 0.009 | 0.648 ± 0.035 | 0.619 ± 0.025 | 0.438 ± 0.016 | 0.705 ± 0.025 |
| Transformer | 0.329 ± 0.068 | 0.270 ± 0.030 | 0.263 ± 0.028 | 0.181 ± 0.033 | 0.330 ± 0.044 |
| FlatMlp | 0.291 ± 0.020 | 0.320 ± 0.042 | 0.206 ± 0.023 | 0.144 ± 0.014 | 0.245 ± 0.021 |
| *AUROC* | | | | | |
| MotifCnn | 0.895 ± 0.004 | 0.919 ± 0.003 | 0.945 ± 0.002 | 0.831 ± 0.005 | 0.955 ± 0.003 |
| DilatedResCnn | 0.882 ± 0.011 | 0.907 ± 0.009 | 0.944 ± 0.008 | 0.824 ± 0.016 | 0.946 ± 0.008 |
| BiLSTM | 0.899 ± 0.004 | 0.921 ± 0.001 | 0.886 ± 0.004 | 0.795 ± 0.009 | 0.928 ± 0.005 |
| BiGRU | 0.894 ± 0.008 | 0.912 ± 0.002 | 0.894 ± 0.009 | 0.795 ± 0.013 | 0.927 ± 0.006 |
| Conv stem + Transformer | 0.898 ± 0.003 | 0.919 ± 0.008 | 0.896 ± 0.007 | 0.805 ± 0.002 | 0.942 ± 0.003 |
| Transformer | 0.726 ± 0.040 | 0.694 ± 0.011 | 0.684 ± 0.012 | 0.616 ± 0.025 | 0.720 ± 0.027 |
| FlatMlp | 0.711 ± 0.007 | 0.719 ± 0.024 | 0.646 ± 0.008 | 0.602 ± 0.004 | 0.691 ± 0.009 |

**Table S4.** TOMTOM analysis of the GUE datasets' motifs. TOMTOM version 5.5.9 was run on the JASPAR 2026 CORE vertebrates non-redundant collection using tomtom -no-ssc -oc . -verbosity 1 -min-overlap 5 -dist pearson -evalue -thresh 10.0 all.meme db/JASPAR/JASPAR2026_CORE_vertebrates_non-redundant.meme. We count a motif family as learned by a seed when one of that seed's filters matches a profile of the family at $q < 0.05$. The cells represent the number of seeds (out of 5) where the family is learned / the median of the five per-seed best q-values / the number of filters at $q < 0.05$ over the five seeds; "-" means no seed reached $q < 0.05$; the families are annotated according to JASPAR, except the MYC/MAX network, which groups MAX, MYC, MYCN, MAX::MYC, MNT and MXI1; all 960 filters of each dataset (5 seeds x 192).

| **Family** | **Pol II (0)** | **Pol II (1)** | **PAX5 (2)** | **KAP1 (3)** | **MXI1 (4)** |
|---|---|---|---|---|---|
| Paired domain only | 1 / 0.35 / 1 | 0 / - / 0 | 5 / $3.2\times10^{-9}$ / 81 | 0 / - / 0 | 0 / - / 0 |
| RFX-related factors | 0 / - / 0 | 0 / - / 0 | 0 / - / 0 | 1 / 0.55 / 1 | 5 / $3.1\times10^{-7}$ / 47 |
| MYC/MAX network (MAX, MYC, MYCN, MAX::MYC, MNT, MXI1) | 1 / 0.063 / 1 | 4 / 0.028 / 5 | 0 / - / 0 | 0 / - / 0 | 5 / $2.4\times10^{-4}$ / 57 |
| Fos-related | 0 / - / 0 | 0 / - / 0 | 3 / 0.020 / 4 | 5 / $2.1\times10^{-5}$ / 55 | 0 / - / 0 |

| Family | Pol II (0) | Pol II (1) | PAX5 (2) | KAP1 (3) | MXI1 (4) |
|---|---|---|---|---|---|
| Jun-related | 0 / - / 0 | 0 / - / 0 | 3 / 0.020 / 4 | 5 / $4.5\times10^{-5}$ / 58 | 0 / - / 0 |
| Three-zinc finger Kruppel-related | 5 / $3.8\times10^{-3}$ / 30 | 5 / $4.0\times10^{-3}$ / 26 | 5 / $5.6\times10^{-4}$ / 33 | 3 / 0.020 / 4 | 4 / 0.011 / 8 |
| Ets-related | 5 / $3.8\times10^{-3}$ / 18 | 5 / $3.0\times10^{-3}$ / 15 | 5 / $1.3\times10^{-4}$ / 39 | 5 / 0.011 / 13 | 5 / $5.4\times10^{-3}$ / 10 |
| More than 3 adjacent zinc fingers | 5 / $5.2\times10^{-4}$ / 89 | 5 / $1.4\times10^{-3}$ / 111 | 5 / $6.1\times10^{-7}$ / 103 | 5 / $1.6\times10^{-3}$ / 33 | 5 / $3.2\times10^{-6}$ / 58 |

### Note S1. Use of AI tools

F.d.S. used Claude Opus 5.5 (Anthropic), through Claude Code, an agentic coding tool that handles file manipulation, coding and approved command invocations, in October 2026. The agentic contribution is found in PR #2 to #6 in the GitHub repository and covers GPU and buffer fixes, the per-epoch callback with ROC/PR curves, the MEME motif export, the crate rename with the removal of the graph models, and the evaluation harness. All changes were manually reviewed, tested and committed.

The agent was used to run the experiments (command invocation, monitoring and running the statistics scripts); every evaluation result comes from executed code and is confirmed by a complete re-run on each device. This includes the TOMTOM run on the five seeds (960 filters per dataset), where it was used to gather all required data and software to match G.S.'s exact command, writing the analysis script, checks, table outputs and iterative analysis (spotting groups etc.).

The agent was also used to search the literature, which found Karpinsky *et al.* 2026, BEND, Tang *et al.* 2025 and DeepBioP.

For his final contribution, F.d.S. used AI to compile the results, numbers and sources, and wrote the full text on his own. The text was then checked with AI for consistency, grammar and spelling, and each correction was reviewed, accepted or rejected manually.

GS wrote the initial text and then used ChatGPT for the consistency, grammar and spelling mistakes. GS coded flodl-bio without any ChatGPT support. After the write of the code, GS checked some of the implemented algorithms and bindings and code that can they be improved for the faster execution and using additional features.

### Note S2. Identity of the GUE TF-binding datasets

The five datasets were identified based on the ENCODE Project Consortium et al. An integrated encyclopedia of DNA elements in the human genome. Nature, 489 (7414):57, 2012, and Zhou et al. (2024) for comparative benchmark.

| GUE dataset | ENCODE experiment | Factor | Cell line | Train / Dev / Test |
|---|---|---|---|---|
| 0 | wgEncodeEH000552 | RNA polymerase II | HUVEC | 32,378 / 1,000 / 1,000 |
| 1 | wgEncodeEH000606 | RNA polymerase II | Progenitor fibroblasts (ProgFib) | 30,672 / 1,000 / 1,000 |
| 2 | wgEncodeEH001546 | PAX5 | GM12892 | 19,000 / 1,000 / 1,000 |
| 3 | wgEncodeEH001776 | KAP1 (TRIM28) | U2OS | 27,294 / 1,000 / 1,000 |
| 4 | wgEncodeEH002829 | MXI1 | H1-hESC | 19,000 / 1,000 / 1,000 |
